# ADHD Symptoms are Associated with the Modular Structure of Intrinsic Brain Networks in a Representative Sample of Healthy Adults

**DOI:** 10.1101/505891

**Authors:** Kirsten Hilger, Christian J. Fiebach

## Abstract

Attention-deficit/hyperactivity disorder (ADHD) is one of the most common neurodevelopmental disorders with significant and often lifelong effects on social, emotional, and cognitive functioning. Influential neurocognitive models of ADHD link behavioral symptoms to altered connections between and within functional brain networks. Here, we investigate whether network-based theories of ADHD can be generalized to understanding variations in ADHD-related behaviors within the normal (i.e., clinically unaffected) adult population. In a large and representative sample, self-rated presence of ADHD symptoms varied widely; only eight out of 291 participants scored in the clinical range. Subject-specific brain-network graphs were modeled from functional MRI resting-state data and revealed significant associations between (non-clinical) ADHD symptoms and region-specific profiles of between-module and within-module connectivity. Effects were located in brain regions associated with multiple neuronal systems including the default-mode network, the salience network, and the central executive system. Our results are consistent with network perspectives of ADHD and provide further evidence for the relevance of an appropriate information transfer between task-negative (default-mode) and task-positive brain regions. More generally, our findings support a dimensional conceptualization of ADHD and contribute to a growing understanding of cognition as an emerging property of functional brain networks.

**Author Summary:** Neurocognitive models of ADHD link behavioral symptoms to altered connections between and within functional brain networks. We investigate whether these network-based theories of ADHD can be generalized to ADHD-related behaviors within the normal adult population. Subject-specific brain graphs were modeled from functional MRI resting-state data of a large and representative sample (*N* = 291). Significant associations between ADHD-related behaviors and region-specific profiles of between-module and within-module connectivity were observed in brain regions associated with multiple functional systems including the default-mode network, the salience network, and the central executive system. Our results support a dimensional conceptualization of ADHD and enforce network-based models of ADHD by providing further evidence for the relevance of an appropriate information transfer between task-negative (default-mode) and task-positive brain regions.

## Introduction

Attention-deficit/hyperactivity disorder (ADHD) is one of the most commonly diagnosed neurodevelopmental disorders with a world-wide prevalence of ~ 5.3% (Polanczyk et al., 2007). Affected patients suffer from symptoms of inattention, impulsivity, and hyperactivity. Although symptoms typically start in childhood, ~ 30-50% of patients are also affected during adult life (Balint et al., 2008), showing persistent problems in social functioning, lower academic success, and higher risk for psychiatric problems (Bussing et al., 2010; Fischer et al., 1990). ADHD has long been treated as categorical concept ignoring considerable symptom variability across (Mostert et al., 2015) and within persons over time (Biederman et al., 2000). In line with a more dimensional conceptualization of ADHD (Marcus et al., 2012), recent research however demonstrates that even non-clinical variations in ADHD symptoms significantly impact cognitive functioning and psychological wellbeing (Brown & Casey, 2016; Groen et al., 2018).

Neuroimaging studies identified associations between ADHD and a wide range of variations in brain structure and function. Reductions in gray matter volume are frequent and most consistently found in prefrontal regions, basal ganglia, and cerebellum (Frodl & Skokauskas, 2012; Konrad et al., 2018). ADHD-related reductions in structural brain connections were observed in cortico-striato-thalamico-cortical loops (Cortese et al., 2013; Konrad et al., 2010), corpus callosum (Pastura et al., 2016; van Ewijk et al., 2014), and the cerebellar peduncles (Ashtari et al., 2005; Nagel et al., 2011). Functional neuroimaging (see Cortese et al., 2012; McCarthy et al., 2014; Norman et al., 2016 for meta-analyses) additionally indicates altered patterns of neural activation during different cognitive tasks, most prominently reduced activation in task-positive regions (executive control network, ventral attention/salience network, striatum; Seeley et al., 2007) and lower levels of task-related deactivation in task-negative regions (default-mode network; Raichle et al., 2001). Investigations of task-free (i.e., resting-state) fMRI have demonstrated disturbed functional connectivity patterns (Rubia, 2018; Konrad et al., 2018), which have been proposed as an intrinsic neural characteristic of ADHD (Castellanos & Aoki, 2016).

Current neurocognitive models of ADHD focus on altered connectivity patterns between functional brain networks: The *default-mode interference hypothesis* (Sonuga-Barke & Castellanos, 2007) postulates that ADHD-related fluctuations and variability in attention and cognition (Castellanos et al., 2005) result from an inadequate regulation of the default-mode network by task-positive networks – which according to this model increases the likelihood of spontaneous and distracting intrusions of introspective thought into ongoing cognitive processes. Empirical support comes from studies reporting increased interactions (decreased anti-correlations) between the default-mode and task-positive networks (Sun et al., 2012; Mills et al., 2018; Mowinckel et al., 2017) as well as systematic hyperactivation in defaultmode regions during cognitive tasks (Cortese et al., 2012; see also the *systems-neuroscience model* of ADHD proposed by these authors). Recent studies broadened the focus towards three-network models of ADHD, proposing that stronger interactions between salience and default-mode network relative to weaker interactions between salience and central executive network reflect a deficient ability of the salience network to adaptively switch between central executive and default-mode network in response to current task demands (Choi et al., 2013).

Graph-theoretical network analysis has emerged as a valuable method for studying network properties of the human brain (Sporns, 2011a,b). Brain networks can be partitioned into subnetworks (communities/modules) that share topological properties and supposedly fulfill specific cognitive or behavioral functions (Sporns & Betzel, 2016). Taking into account this modular structure of the human brain makes it possible to examine region-specific interactions between and within different networks in a quantifiable manner – and to test neurocognitive models of ADHD. First graph-theoretical investigations of ADHD-related network organization (Lin et al., 2014; Barttfeld et al., 2014), however, studied modularity only at a whole-brain level, i.e., as global property of the entire brain. Accordingly, this work cannot inform about altered connection patterns between or within different networks/modules as postulated by neurocognitive theories of ADHD. Here, we aim at relating graph-theoretical network analysis more directly to network models of ADHD by analyzing two local graph-theoretical measures that provide complementary information about a brain region’s connections within and between different modules.

Finally, it is noteworthy that empirical support for network models of ADHD so far primarily comes from categorical studies of ADHD, i.e., group-level comparisons between patients and controls (Sun et al., 2012; Choi et al., 2013). As outlined above, this approach ignores recent advances towards a more dimensional understanding of ADHD (Marcus et al., 2012). As of now, it accordingly remains unclear whether network models of ADHD are valid only for clinically affected persons or whether they may also inform more generally about mechanisms linking between-person variation in brain network organization to variation in cognition. To address this question, we applied graph-theoretical modularity analyses to a large and representative sample of *N* = 291 adults.

## Methods

The current study was conducted with data acquired at the Nathan S. Kline Institute for Psychiatric Research and online distributed as part of the 1000 Functional Connectomes Project INDI (Enhanced NKI Rockland Sample, Nooner et al., 2012, http://fcon_1000.projects.nitrc.org/indi/enhanced/). Experimental procedures were approved by the Nathan S. Kline Institute Institutional Review Board (#239708), and informed written consent was obtained from all participants. Note that data acquisition, preprocessing, and graph-theoretical analyses are to a large degree identical with a previous publication from our research group, which however focused on a different outcome measure (Hilger et al., 2017b). The code used in the current study has been deposited on github at https://github.com/KirstenHilger/ADHD-Modularity (https://doi.org/10.5281/zenodo.2574588).

### Participants

301 participants with complete phenotypical and neuroimaging data were selected from the Enhanced NKI Rockland sample. Two participants were excluded due to medication with methylphenidate, which can alter neural activation related to ADHD symptoms (e.g., Shafritz et al., 2004); eight participants were excluded due to high in-scanner motion, i.e., mean framewise displacement (*FD*) > 0.2 mm. Thus, our final sample comprises 291 participants (18-60 years, *M* = 39.34, *SD* = 13.80; 189 females; handedness: 251 right, 21 left, 19 ambidextrous). ADHD symptoms were assessed with the Conners’ Adult ADHD Rating Scales (Conners et al., 1999; Self Report, Short Version/CAARS-S:S), from which four subscale scores (Inattention/Memory Problems, Hyperactivity/Restlessness, Impulsivity/Emotional Lability, and Problems with Self-Concept) as well as the total Index of ADHD symptoms were computed. The ADHD Index was used as variable of interest in all graph analyses. Potential differences in brain network organization due to different levels of intelligence (e.g., Hilger et al., 2017a; van den Heuvel et al., 2009) were controlled by using the Full Scale Intelligence Quotient (FSIQ; Wechsler Abbreviated Scale of Intelligence, WASI, Wechsler, 1999; range 68 to 135, *M* = 99.22, *SD* =12.50) as covariate of no interest.

### fMRI Data Acquisition

Resting-state fMRI data were acquired on a 3 Tesla whole-body MRI scanner (MAGNETOM Trio Tim, Siemens, Erlangen, Germany). A T2*-weighted BOLD-sensitive gradient-echo EPI sequence was measured with the following parameter: 38 transversal axial slices of 3mm thickness, 120 volumes, field of view [FOV] 216×216mm, repetition time [TR] 2500ms, echo time [TE] 30ms, flip angle 80°, voxel size 3×3×3mm, acquisition time 5.05min. Further, three-dimensional high-resolution anatomical scans were obtained for coregistration with a sagittal T1-weighted, Magnetization Prepared-Rapid Gradient Echo sequence (176 sagittal slices, FOV 250×50mm, TR 1900ms, TE 2.5ms; flip angle 9°, voxel size 1×1×1mm, acquisition time 4.18min).

### fMRI Data Preprocessing

Preprocessing of neuroimaging data was conducted using the freely available software packages AFNI (http://afni.nimh.nih.gov/afni) and FSL (http://www.fmrib.ox.ac.uk/fsl/). The first four EPI volumes were discarded to allow for signal equilibration. Next steps comprised slice-time correction, three-dimensional motion correction, time-series despiking, and spatial smoothing (6mm FWHM Gaussian kernel). Four-dimensional mean-based intensity normalization was performed and data were temporally filtered with a bandpass filter of 0.005-0.1Hz. Linear and quadratic trends were removed and all individual EPI volumes were normalized to the MNI152 template (3×3×3 mm resolution) using nonlinear transformations and each individual’s anatomical scan. Finally, nine nuisance signals were regressed out, i.e., six motion parameters (rigid body transformation) as well as regressors for cerebrospinal fluid (intra-axial), white matter, and global mean signal that were calculated by averaging (AFNI, 3dmaskave) voxel-wise BOLD time series within subject-specific masks resulting from FSL’s automatic segmentation (FAST) of the anatomical image. Framewise displacement (*FD*) was calculated on the basis of the six motion parameters indicating translation/rotation in three directions between two consecutive volumes, *FD_i_* = |*Δd_ix_*| + |*Δd_iy_*| + |*Δd_iz_*| + |*Δα_i_*| + |*Δβ_i_*| + |*Δγ_i_*| (Power et al., 2012); subjects with mean *FD* > 0.2 mm were excluded (*N* = 8; see above). In-scanner motion (mean *FD*) was not significantly related to the variable of interest, i.e., ADHD Index (*r* = -.05, *p* = .409). Nevertheless, to further minimize potential remaining influences of head motion on the observed effects, we added mean *FD* as control variable in all individual difference analyses. For subsequent graph analyses, data were downsampled by factor two, resulting in individual maps of 6×6×6mm resolution. The preprocessing scripts used in the current study were released by the 1000 Functional Connectomes Project and are available for download at http://www.nitrc.org/projects/fcon_1000.

### Graph-theoretical Analyses of Intrinsic Connectivity

Individual brain graphs were constructed on the base of all 5,411 gray matter voxels in the brain, which served as nodes for the respective graphs. Network edges were modeled between nodes showing high positive correlations of BOLD signal time series. Edges between physically close nodes (< 20 mm) were excluded, as they may result from motion artifacts and spuriously high correlations induced by shared nonbiological signals (Power et al, 2011). Community detection and the subsequent graph metrics were calculated for five separate graphs defined by five proportional thresholds (representing the top 10%, 15%, 20%, 25%, and 30% of strongest edges, i.e., highest correlations), which also excluded all negative network edges (Murphy et al., 2009). The subject-specific averages of graph metrics across these five thresholds were used in all following analyses (see also Hilger et al., 2017b). Finally, all graphs were binarized (as recommended for individual difference analyses; van Wijk et al., 2010).

### Global Modularity

To study the modular organization of intrinsic brain networks, each individual network graph was parcellated into several functionally distant communities or modules. To this end, we applied the standard Louvain algorithm (Blondel et al., 2008), which finds the optimal modular partition in an iterative procedure by maximizing the *global modularity Q* (Newman & Girwan, 2004):

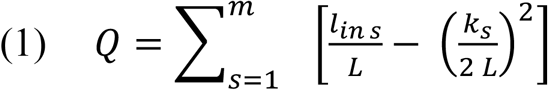

Here, *m* represents the number of modules, *l_in s_* is the number of edges inside module *s, L* reflects the total number of edges in the network, and *k_s_* represents the total degree of the nodes in module *s*. Thus, the first term of formula (1) represents the actual fraction of within-module edges relative to all edges in the network, whereas the second term represents the expected fraction of within-module edges. When the first term (actual within-module edges) is much higher than the second term (expected within-module edges), many more edges are present inside module s than expected by chance. In this case, the *global modularity Q*, which results from summing up these differences over all modules *m* in the network, increases. Usually, modularity values above 0.3 are taken as indicator of a network’s modular organization (Fortunato & Barthélemy, 2007). In addition to *global modularity Q*, we calculated three further whole-brain measures of modular network organization for the final module partition of each participant, i.e., *number of modules, average module size*, and *variability in module size*.

### Node-specific modularity measures

Node-specific analyses of network modularity were conducted using *participation coefficient p_i_* and *within-module degree z_i_*. The *participation coefficient p_i_* assesses the diversity of each node’s connections across all modules in the brain (Bertolero et al., 2017) and is defined as:

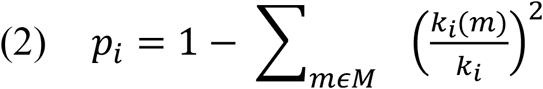

Here, *k_i_* is the degree of node *i* and thus represents the number of edges directly attached to node *i. k_i_*(*m*) refers to the subset of edges linking node *i* to other nodes within the same module *m* (Rubinov & Sporns, 2010; Guimerà & Amaral, 2005). The *participation coefficient p_i_* is 1 when a node is equally connected to all modules within the network, while it is 0 when all of its connections are to one single module (Bertolero et al., 2015, 2017, 2018; Sporns & Betzel, 2016). *Within-module degree z_i_*, in contrast, represents within-module connectivity and is defined as:

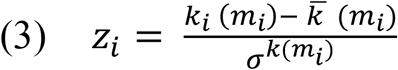

Here, *m_i_* is the module of node *i. k_i_*(*m_i_*) represents the number of connections within the node’s own module (i.e., the within-module degree of node *i*), and 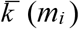 and *σ*^*k*(*mi*)^ are the mean and standard deviation of the within-module degree distribution of module *m_i_* (Guimerà & Amaral, 2005). Nodes that are highly connected to nodes within their own module receive positive values of *within-module degree z_i_*, whereas nodes with low levels of connectivity within their own module are characterized by negative values (Sporns & Betzel, 2016). All graph-theoretical network analyses were conducted in python using the open-source software *network-tools* (Ekman & Linssen, 2015).

### Node-type analysis

Functional cartography (Guimerà & Amaral, 2005) relies on the above-described node-specific metrics (i.e., *participation coefficient p_i_, within-module degree z_i_*) and can be used to assign each network node into one of seven different classes, which are in turn characteristic for the node’s role in within- and between-module communication (see Figure 1C). As suggested in the original work of Guimerà and Amaral (2005) and used in previous studies (e.g., Sporns et al., 2011), nodes with *within-module degree z_i_* ≥ 1 were classified as hubs (17,86% of all nodes) and nodes with *z_i_* < 1 were classified as non-hubs. On the basis of the *participation coefficient p_i_*, non-hubs were further classified as *ultra-peripheral* (*p_i_* ≤ 0.05), *peripheral* (0.05 < *p_i_* ≤ 0.62), *non-hub connector* (0.62 <*p_i_* ≤ 0.80), or *non-hub kinless nodes* (*p_i_* > 0.80), whereas hubs were classified as *provincial* (*p_i_*. ≤ 0.30), *connector* (0.30 < *p_i_*. ≤ 0.75), or *kinless hubs* (*p_i_* > 0.75).

**Figure 1.**
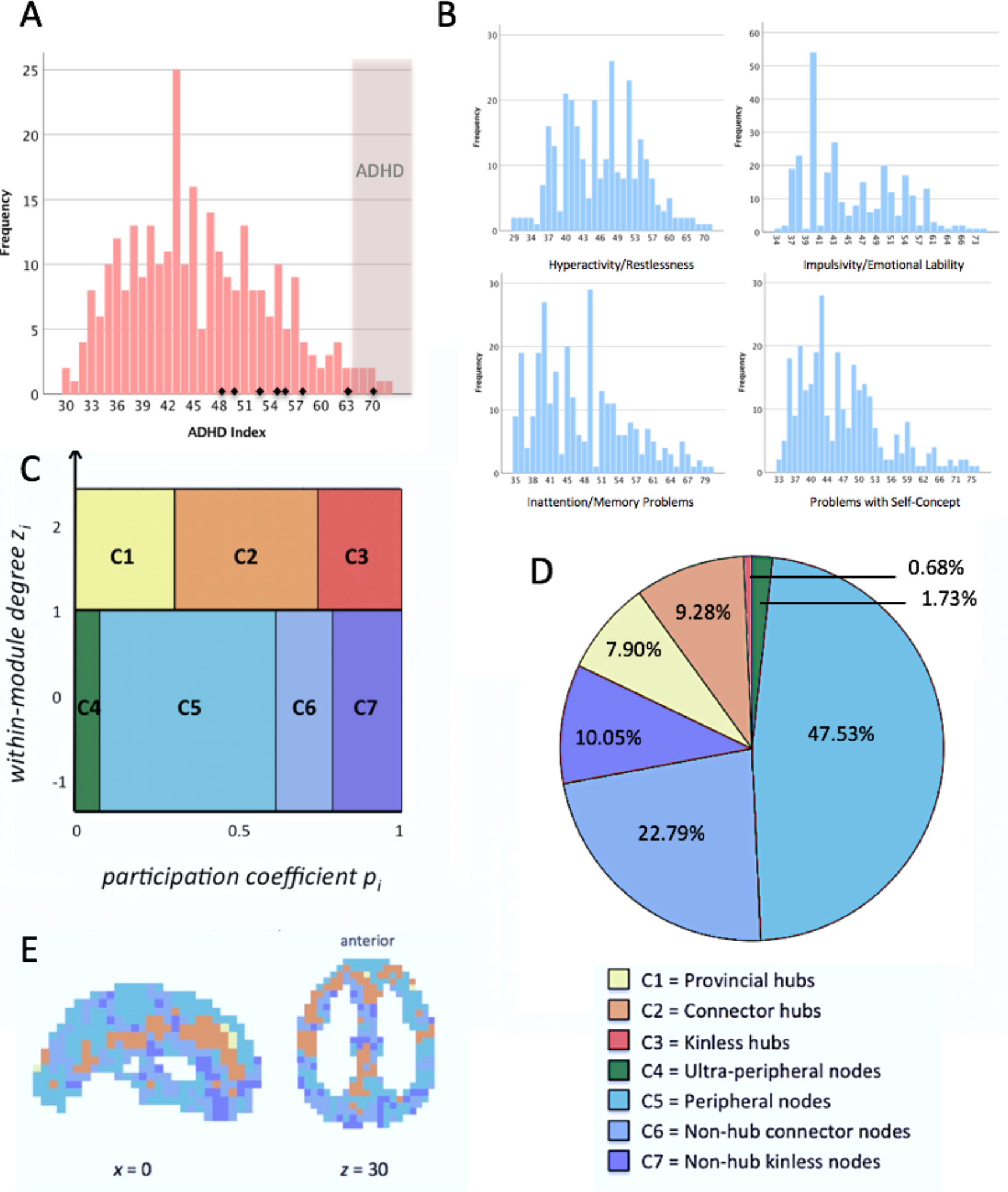
Frequency plots of CAARS subscales and illustration of node-type analysis. **(A)** Frequency histogram of Conners’ ADHD Index *t*-scores. Values > 65 describe the 95-98^th^ percentile, are interpreted as “much above average”, and suggest the presence of ADHD (Conners et al., 1999). The respective area is depicted in light red. Subjects with clinical ADHD diagnosis are illustrated as small black diamonds in the histogram. **(B)** Frequency histograms of t-scores of CAARS subscales. **(C)** Seven node types defined as a function of their profile of between-module and within-module connectivity, i.e., *participation coefficient p_i_* and *within-module degree z_i_*. Adapted from Guimerà and Amaral (2005; functional cartography; see Methods). **(D)** Proportions of node types within the whole brain and across all subjects. The proportions of node types were calculated for each subject separately and averaged across all participants afterwards. **(E)** Anatomical distribution of node types within the whole brain, depicted here for exemplary purposes for one subject. Hubs (*z_i_* > 1) are illustrated in warm colors (yellow to red), non-hub nodes (*z_i_* ≤ 1) are illustrated in cool colors (green to blue). Subject-specific values of *participation coefficient p_i_* and *within-module degree z_i_* as well as the respective proportions of node types within the whole brain were calculated for proportionally thresholded and binarized graphs (five different cut-offs, i.e., the top 10%, 15%, 20%, 25%, or 30% of strongest edges were used to model the graph). Individual node-type proportions were calculated by averaging across the five thresholds and averaged across all subjects afterwards. The *x*- and *z*-coordinates represent coordinates of the Montreal Neurological Institute template brain (MNI152).

### ADHD symptoms and differences in modular network organization

The primary aim of the present study was to examine whether or not individual differences in the strength of ADHD-related behavior, in a non-clinical sample, are associated with individual differences in modular brain network organization. To this end, partial correlations were calculated between the Conners’ ADHD Index and global measures of modular brain organization, i.e., *global modularity Q, number of modules, average module size*, and *variability in module size*, as well as whole-brain proportions of the seven node types. In these analyses, we controlled for effects of age, sex, handedness, FSIQ, and mean *FD*, and excluded outliers, i.e., subjects with values > 3 *SD* above/below the mean of the respective variable of interest. Statistical significance was accepted at *p* < .05, however correcting for multiple comparisons using Bonferroni correction, resulting in *p*-thresholds of .013 for global modularity measures (4 analyses) and .007 for node-type proportions (7 analyses). To quantify the evidence in favor of the null hypothesis (i.e., absence of an association) for nonsignificant correlation results, we calculated Bayes Factors (BF_01_; Jeffreys, 1961; Wetzels & Wagenmakers, 2012) using Bayesian linear regression and the default prior (Rouder & Morey, 2012) as implemented in JASP (Version 0.8.6; https://jasp-stats.org). Substantial evidence for the null was accepted at BF_01_ > 3 (Jeffreys, 1961).

Finally, associations between Conners’ ADHD Index and node-specific (i.e., voxel-wise) measures of modular network organization (i.e., *within-module degree z_i_, participation coefficient p_i_*) were examined using regression models in the Statistic Parametric Mapping software (Welcome Department of Imaging Neuroscience, London, UK), again controlling for age, sex, handedness, intelligence (FSIQ), and mean *FD*. The resultant *p*-values were FWE-corrected for multiple comparisons with Monte Carlo-based cluster-level thresholding as implemented in AFNI (Forman et al., 1995), whereby an overall threshold of *p* < .05 was achieved by combining a voxel threshold of *p* < .005 with a cluster-based extent threshold of *k* > 26 voxels (3dClustSim; AFNI version August 2016; voxel size 3×3×3mm; 10,000 permutations; Ward, 2000).

## Results

### ADHD-related behavior

A descriptive characterization of the distribution of self-rated ADHD symptoms as assessed with the Conners’ Adult ADHD Rating Scale (CAARS) is presented in Figure 1 A,B (see also Table 1A). As expected for a representative adult sample, the distribution is positively (i.e., right) skewed and the majority of participants exhibited ADHD Index values clearly below the threshold for ADHD diagnosis, i.e., *t*-scores < 65 (Conners et al., 1999). Nevertheless, we observed considerable variation between participants in the global ADHD Index (Figure 1A) and its four subscales, i.e., inattention, impulsivity, hyperactivity, problems with self-concept (Figure 1B), suggesting that the CAARS is well suited for describing non-clinical between-person variations in ADHD-related behavior. Note that although eight participants reported a clinical diagnosis of ADHD, only one of them fulfilled the CAARS’ ADHD criteria. Nevertheless, all participants with clinical diagnosis fell within the upper half of the distribution (Figure 1A).

**Table 1.** Descriptive statistics of the CAARS scales and global modularity measures

|  | *M* | *SD* | *Median* | *Min* | *Max* |
| --- | --- | --- | --- | --- | --- |
| *ADHD symptoms* |  |  |  |  |  |
| ADHD Index | 8.36 | 5.13 | 8.00 | 0 | 28 |
| inattention | 3.82 | 3.83 | 4.00 | 0 | 13 |
| hyperactivity | 4.42 | 2.65 | 4.00 | 0 | 12 |
| impulsivity | 3.07 | 2.32 | 2.00 | 0 | 12 |
| self-concept problems | 4.24 | 3.22 | 4.00 | 0 | 15 |
| *Whole-brain modularity measures* |  |  |  |  |  |
| global modularity | .37 | .03 |  | .30 | .48 |
| number of modules | 3.54 | 0.33 |  | 2.80 | 4.80 |
| average module size | 1572.35 | 137.98 |  | 1149.01 | 2073.07 |
| variability in module size | 371.04 | 161.55 |  | 94.49 | 1105.95 |

*M*, mean; *SD*, standard deviation; *Min*, minimum value observed across all participants; *Max*, maximum value observed across all participants. Statistics for the whole-brain modularity measures refer to subject-specific values after averaging across all graph-defining thresholds, i.e., 10%, 15%, 20%, 25%, 30%. The variables *average module size* and *variability in module size* were measures in nodes.

### Modular brain network organization

Table 1B reports descriptive statistics for global characteristics of modular network organization, i.e., *global modularity Q, number of modules, average module size*, and *variability in module size*. Mean values for *global modularity Q* were all greater than .3, indicating a clear modular organization in all participants (Fortunato & Barthélemy, 2007). The proportions of functionally different node types (Figure 1C,D) are likely to influence the global information flow (van den Heuvel & Spons, 2013), and are thus also considered as global properties of modular network organization. As we would assume for a network whose overall organization is characterized by substantial modularity and small-worldness (Gallos et al., 2012; Sporns & Betzel, 2016), only a minority of nodes were characterized as hubs (i.e., connector, provincial, or kinless hubs; in total 17.86%) and the most common node types were peripheral nodes and non-hub connector nodes.

For most participants, the anatomical distribution of node types matched observations of previous studies (see e.g., Meunier et al., 2010): For example, connector hubs were localized along the midline and on the borders between anatomically segregated cortices, whereas less important nodes were observed in more peripheral and functionally specialized regions.

Figure 1E visualizes this anatomical distribution for one randomly selected participant (see also Hilger et al., 2017b).

The group-average spatial distribution of the two nodal measures, *participation coefficient p_i_* and *within-module degree z_i_*, matched nearly perfectly the distribution we recently published for a slightly larger sub-sample from the same dataset (see Figure 1 in Hilger et al., 2017b), and is therefore not visualized again. Network nodes with high values of *participation coefficient p_i_* were located in medial prefrontal (i.e., anterior and mid-cingulate) cortex, anterior insula, inferior frontal gyrus, superior temporal gyrus, medial temporal structures (amygdala, hippocampus), inferior parietal lobule, posterior cingulate cortex/precuneus, and in the thalamus. Nodes with high *within-module degree z_i_* were observed in large parts of medial prefrontal cortex (again including anterior and mid-cingulate cortex), supplementary motor area, lateral superior and middle frontal gyri, anterior insula, postcentral gyrus, temporo-parietal junction, posterior cingulate cortex/precuneus, middle occipital/lingual gyrus, and in the cuneus.

### Local but not global measures of modular brain network organization are associated with ADHD symptoms

Neither global measures of modular organization (*global modularity Q, number of modules, average module size*, and *variability in module size*) nor the whole-brain proportions of node types were significantly associated with Conners’ ADHD Index (Table 2). Bayes Factors exceeded 3 in only two out of 11 analyses, indicating that despite the relatively large sample of almost 300 participants, further evidence would be needed to achieve robust support against an association between ADHD symptoms and global modularity measures.

**Table 2.** ADHD symptoms and global modularity measures

| | $r_{part.}$ | $p_{part.}$ | BF<sub>01</sub>-Reg. |
| --- | --- | --- | --- |
| *Whole-brain modularity measures* |  |  |  |
| global modularity | .11 | .056 | 0.62 |
| number of modules | -.09 | .150 | 1.36 |
| average module size | .08 | .213 | 1.95 |
| variability in module size | -.02 | .781 | 3.47 |
| *Whole-brain proportions of node types* |  |  |  |
| ultra-peripheral nodes | .01 | .813 | 3.78 |
| peripheral nodes | .06 | .363 | 2.29 |
| non-hub connector nodes | -.07 | .229 | 2.30 |
| non-hub kinless nodes | -.10 | .103 | 0.98 |
| provincial hubs | .09 | .148 | 1.33 |
| connector hubs | -.07 | .252 | 2.56 |
| kinless hubs | -.07 | .255 | 2.58 |

$r_{part.}$ , Pearson's correlation coefficient for the partial correlation controlling for effects of age, sex, handedness, and FSIQ; $p_{part.}$ , $p$ -value of significance for the partial-correlation; $BF_{01}$ -Reg., Bayes Factor in favor of the null hypothesis (i.e., absence of correlation). Bayes Factors were calculated for linear regression models predicting ADHD Index values by the respective whole-brain measure of modular network organization or whole-brain proportions of node types, respectively, while effects of age, sex, handedness, mean $FD$ , and FSIQ were controlled.

However, we observed significant associations between Conners’ ADHD Index and node-specific characteristics (Tables 3–5; Figures 2,3). Thus, although individual variation in ADHD-related behaviors did not relate to global properties of modularity, there was a systematic association with the embedding of specific brain regions into the communication between and within different modules. Positive associations between ADHD Index and *participation coefficient p_i_* were observed in five clusters, i.e., in left and right posterior insula (extending laterally into the superior temporal gyri), anterior cingulate cortex, posterior-medial superior frontal gyrus (supplementary motor area), as well as in the left inferior parietal lobe (primarily supramarginal gyrus). Negative associations were observed in eight clusters, including anterior cingulate gyrus, right lateral middle frontal gyrus, left supplementary motor area, left posterior fusiform gyrus, right intraparietal sulcus, right posterior cingulate cortex/precuneus, posterior middle temporal gyrus adjacent to the occipital cortex, and in the right inferior parietal lobe (Table 3, Figure 2).

**Figure 2.**
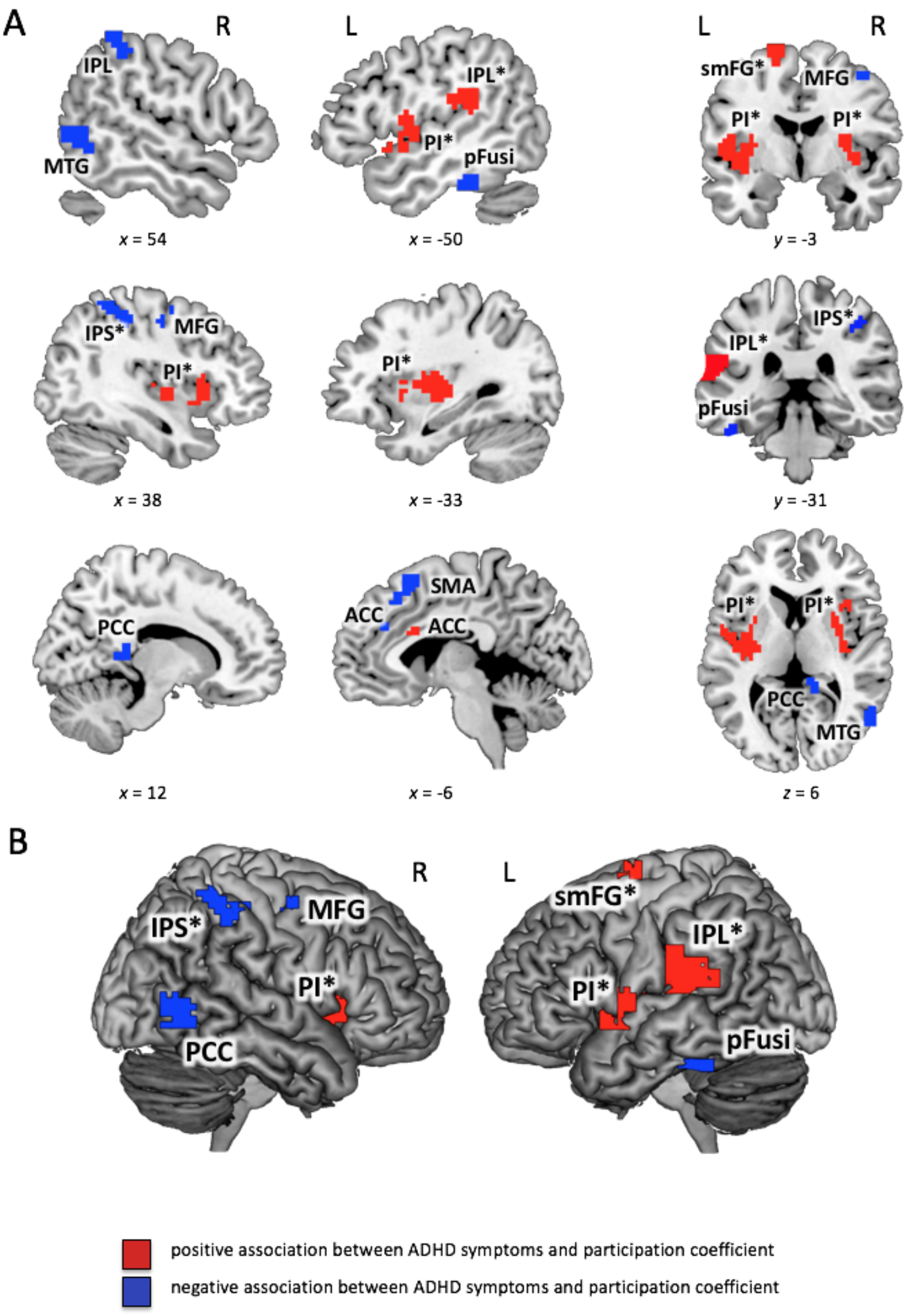
Significant associations between Conners’ ADHD Index and participation coefficient (see also Table 3). *Participation coefficient p_i_* (see Methods for details) was calculated for binarized and proportionally thresholded graphs using five thresholds (graphs were defined by the top 10%, 15%, 20%, 25%, or 30% of strongest edges). Input for analyses were the individual mean maps for *participation coefficient p_i_*, which were calculated by averaging across these five thresholds for each participant separately. Statistic parametric maps of *participation coefficient p_i_* are shown at a voxel-level threshold of *p* < .005 (uncorrected) combined with a cluster-level threshold of *k* > 26 voxels, corresponding to an overall family-wise error corrected threshold of *p* < .05 (see Methods). Clusters with effects in both (between-module and within-module connectivity, i.e., *p_i_* and *z_i_*) are marked with an asterisk (see also Table 5). **(A)** Slice view, the *x*-, *y*-, and *z*-coordinates represent coordinates of the Montreal Neurological Institute template brain (MNI152). **(B)** Render view, projection to the surface of the brain, search depth 12 voxels. PI, posterior insula; IPL, inferior parietal lobe; IPS, intraparietal sulcus; ACC, anterior cingulate cortex; MFG, middle frontal gyrus; SMA, supplementary motor area; pFusi, posterior fusiform gyrus; PCC, posterior cingulate cortex; MTG, middle temporal gyrus; smFG, superior medial frontal gyrus.

**Table 3.** ADHD symptoms and participation coefficient

| Brain Region | BA | Hem | x | y | z | $t_{max}$ | $k$ |
| --- | --- | --- | --- | --- | --- | --- | --- |
| *positive association* |  |  |  |  |  |  |  |
| posterior inula* | 13 | L | -33 | -9 | 9 | 3.78 | 325 |
| posterior inula, putamen* | 13 | R | 33 | 18 | 0 | 3.48 | 207 |
| anterior cingulate cortex | 24 | L | -6 | 18 | 24 | 3.26 | 44 |
| superior medial frontal gyrus* | 6 | L | -18 | -3 | 69 | 3.27 | 38 |
| inferior parietal lobe* | 40 | L | -57 | -36 | 21 | 4.91 | 250 |
| *negative association* |  |  |  |  |  |  |  |
| anterior cingulate cortex | 32, 9 | L | -9 | 36 | 30 | 3.27 | 37 |
| middle frontal gyrus | 6 | R | 33 | -12 | 45 | 3.22 | 53 |
| supplementary motor area | 8, 6 | L | -6 | 18 | 57 | 3.65 | 49 |
| posterior fusiform gyrus | 20, 36 | L | -51 | -36 | -27 | 4.08 | 49 |
| intraparietal sulcus* | 40 | R | 33 | -36 | 48 | 3.33 | 90 |
| posterior cingulate cortex |  | R | 12 | -39 | 9 | 3.71 | 26 |
| middle temporal gyrus | 37, 19 | R | 57 | -63 | 0 | 3.77 | 91 |
| inferior parietal lobe | 40 | R | 57 | -39 | 51 | 3.08 | 36 |

BA, approximate Brodmann's area; Hem, hemisphere; L, left; R, right; regions with significant effects in both measures (between-module and within-module connectivity) are marked with an asterisk and separately listed in Table 5; coordinates refer to the Montreal Neurological Institute template brain (MNI); $t_{max}$ , maximum $t$ statistic in the cluster; $k$ , cluster size in voxels of size 3 x 3 x 3 mm.

**Table 4.** ADHD symptoms and within-module degree

| Brain Region | BA | Hem | x | y | z | $t_{max}$ | $k$ |
| --- | --- | --- | --- | --- | --- | --- | --- |
| *positive association* |  |  |  |  |  |  |  |
| supplementary motor area | 8, 6 | R/L | 0 | 30 | 60 | 3.50 | 49 |
| temporal cortex, amygdala, hippocampus, fusiform gyrus | 38, 20, 28 | R | 27 | 6 | -33 | 5.31 | 577 |
| temporal cortex, amygdala, hippocampus, fusiform gyrus | 20, 38 | L | -33 | -21 | -33 | 4.63 | 606 |
| precentral gyrus, postcentral gyrus, inferior parietal lobe* | 3, 40 | R | 42 | -33 | 51 | 6.28 | 619 |
| precentral gyrus, postcentral gyrus, inferior parietal lobe | 3, 40 | L | -45 | -33 | 51 | 5.22 | 397 |
| paracentral lobule | 6, 4 | L | -12 | -42 | 72 | 3.73 | 171 |
| *negative association* |  |  |  |  |  |  |  |
| medial frontal gyrus | 11, 32, 25 | R | 3 | 30 | -15 | 3.96 | 119 |
| anterior cingulate cortex | 24 | R | 9 | 18 | 24 | 3.70 | 27 |
| insula, putamen, superior temporal gyrus, inferior frontal gyrus, inferior parietal lobule* | 13, 22, 40 | L | -48 | 9 | -3 | 5.23 | 862 |
| insula, putamen, superior temporal gyrus, inferior frontal gyrus, inferior parietal lobule* | 13, 47, 22 | R | 33 | 0 | 15 | 5.54 | 606 |
| superior medial frontal gyrus* | 6, 32, 24 | L | -12 | -3 | 63 | 5.37 | 497 |
| middle temporal gyrus | 21, 20 | R | 63 | -6 | -21 | 3.93 | 43 |
| thalamus |  | R | 15 | -18 | 9 | 4.02 | 29 |
| thalamus |  | L | -28 | -24 | 6 | 3.44 | 30 |
| posterior fusiform gyrus | 37, 19 | L | -36 | -48 | -12 | 4.38 | 187 |
| posterior fusiform gyrus |  | R | 36 | -57 | -6 | 3.27 | 33 |
| posterior cingulate cortex | 30 | L | -24 | -66 | 21 | 3.70 | 47 |
| precuneus | 19, 7, 31 | R/L | 6 | -78 | 33 | 3.79 | 57 |
| inferior occipital gyrus | 18 | R | 27 | -84 | -12 | 4.09 | 79 |

BA, approximate Brodmann's area; Hem, hemisphere; L, left; R, right; regions with significant effects in both measures (between-module and within-module connectivity) are marked with an asterisk and separately listed in Table 5; coordinates refer to the Montreal Neurological Institute template brain (MNI); $t_{max}$ , maximum $t$ statistic in the cluster; $k$ , cluster size in voxels of size 3 x 3 x 3 mm.

**Table 5.** ADHD symptoms and effects in both participation coefficient and within-module degree

| Brain Region | BA | Hem | x | y | z | k |
| --- | --- | --- | --- | --- | --- | --- |
| *positive association with $p_i$ and negative association with $z_i$* |  |  |  |  |  |  |
| posterior insula, superior temporal gyrus, putamen | 13, 22 | L | -57 | 0 | 9 | 237 |
| posterior insula, putamen | 13 | R | 27 | -15 | 9 | 137 |
| superior medial frontal gyrus | 6 | L | -21 | -6 | 69 | 29 |
| inferior parietal lobe | 40 | L | -69 | -39 | 24 | 67 |
| *positive association with $z_i$ and negative association with $p_i$* |  |  |  |  |  |  |
| intraparietal sulcus | 40 | R | 27 | -36 | 45 | 79 |

BA, approximate Brodmann's area; Hem, hemisphere; L, left; R, right; coordinates refer to the Montreal Neurological Institute template brain (MNI); $t_{max}$ , maximum $t$ statistic in the cluster; $k$ , cluster size in voxels of size 3 x 3 x 3 mm.

Concerning within-module connectivity, ADHD Index was positively associated with *within-module degree z_i_* in six clusters of nodes (see Table 4, Figure 3). Two extensive temporal clusters were observed bilaterally, comprising not only lateral temporal cortices but also the amygdalae, hippocampi, and anterior fusiform gyri. Further, two large central clusters were identified that extended across central and postcentral sulci, from precentral and postcentral gyri to the supramarginal gyri and anterior parts of intraparietal sulci. Smaller clusters were observed in supplementary motor area and superior portions of left post- and precentral gyri (paracentral lobule).

**Figure 3.**
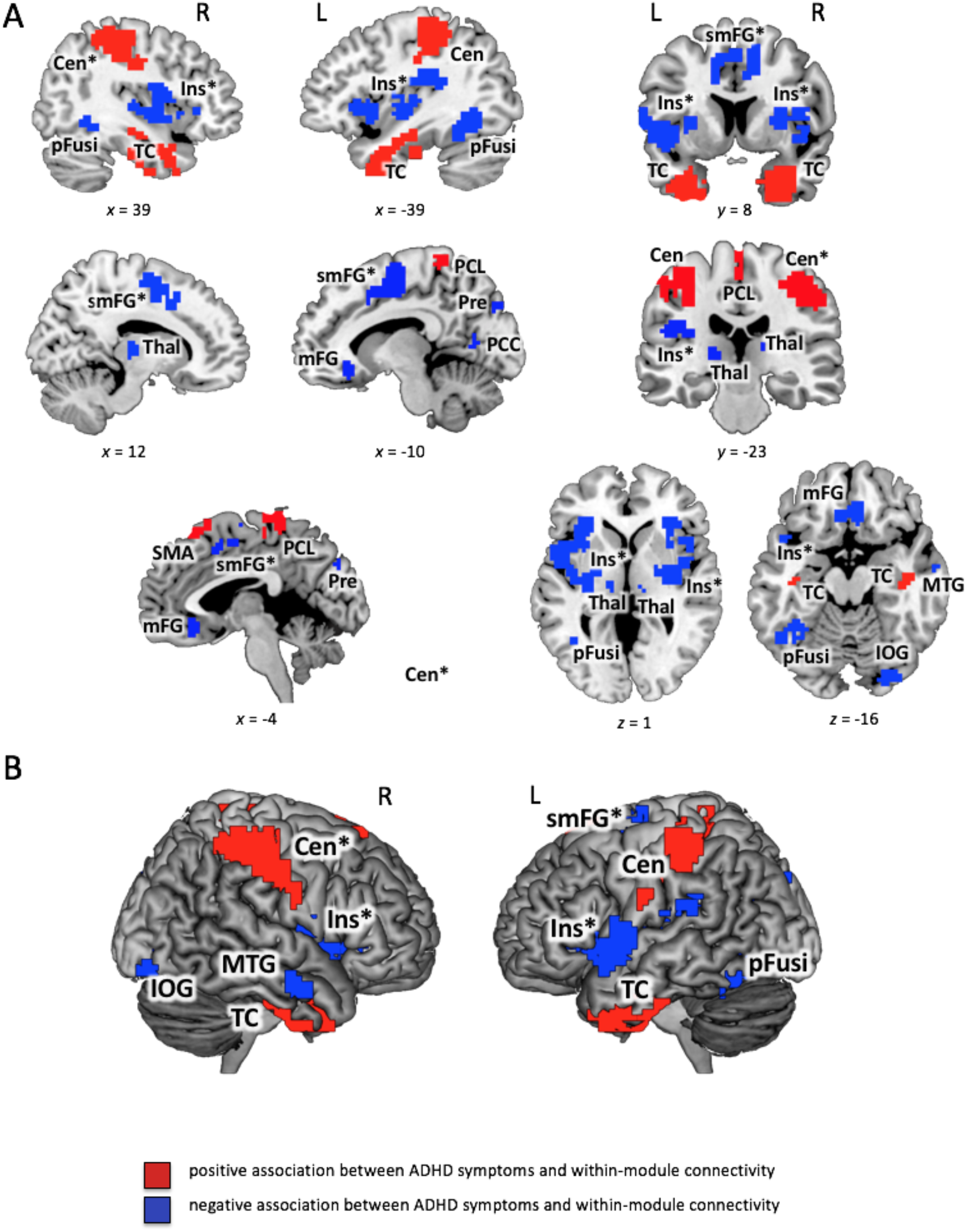
Significant associations between Conners’ ADHD Index and within-module degree (see also Table 4). *Within-module degree z_i_* (see Methods for details) was calculated for binarized and proportionally thresholded graphs using five thresholds (graphs were defined by the top 10%, 15%, 20%, 25%, or 30% of strongest edges). Input for analyses were the individual mean maps for *within-module degree z_i_*, which were calculated by averaging across these five thresholds for each participant separately. Statistic parametric maps of *within-module degree z_i_* are shown at a voxel-level threshold of *p* < .005 (uncorrected) combined with a cluster-level threshold of *k* > 26 voxels, corresponding to an overall family-wise error corrected threshold of *p* < .05 (see Methods). Clusters with effects in both (between-module and within-module connectivity, i.e., *p_i_* and *z_i_*) are marked with an asterisk (see also Table 5). **(A)** Slice view, the *x*-, *y*-, and *z*-coordinates represent coordinates of the Montreal Neurological Institute template brain (MNI152). **(B)** Render view, projection to the surface of the brain, search depth 12 voxel. TC, temporal cluster comprising also amygdala, hippocampus, and parts of fusiform gyrus; Cen, central cluster spreading across central and postcentral sulci from precentral gyri and postcentral gyri to the inferior parietal lobes (comprising supramarginal gyri and anterior parts of intraparietal sulci); PCL, paracentral lobule; mFG, medial frontal gyrus; Ins, insular cluster comprising also parts of putamen, superior temporal gyrus, inferior frontal gyrus and inferior parietal lobe; MTG, middle temporal gyrus; pFusi, posterior fusiform gyrus; Pre, precuneus; IOG, inferior occipital gyrus; smFG, superior medial frontal gyrus; Thal, Thalamus; SMA, supplementary motor area.

Negative associations between ADHD Index and *within-module degree z_i_* were observed in 13 node clusters. Four of those clusters were located medially, in inferior parts of medial frontal gyrus (border to rostral anterior cingulate cortex), in superior parts of postero-medial frontal gyrus, dorsal anterior cingulate cortex, and the precuneus. More laterally located clusters comprised nearly the entire bilateral insulae and reached laterally into the inferior frontal, superior temporal, and inferior parietal lobes (supramarginal gyri), as well as medially into the putamen. Further negatively associated node clusters were observed in right middle and inferior temporal gyri, bilateral thalami, bilateral posterior fusiform gyri, left posterior cingulate cortex, and right inferior occipital cortex. In general, the spatial distribution of significantly associated network nodes showed a surprisingly high degree of inter-hemispheric symmetry for both measures.

In several brain regions, self-rated ADHD-related behaviors were associated with both, i.e., *participation coefficient* and *within-module degree*. These involve nine of the above-described clusters (marked with an asterisk in Figures 2,3; cf. also Table 5). In participants with higher ADHD Index, posterior insulae, left postero-medial superior frontal gyrus, and left inferior parietal lobe showed higher *participation coefficient* along with relatively lower *within-module degree*. The opposite pattern, i.e., lower *participation coefficient* and higher *within-module degree* was observed in the right intraparietal sulcus.

Similar effects were observed for the four subscales of the CAARS (see Supplementary Tables S1 and S2 for associations with global measures).

### Post-hoc analyses

One of the few studies that so far investigated the association between ADHD and graph-theoretical brain network characteristics observed higher *global modularity* in patients with ADHD (Lin et al., 2014) – which is not consistent with our results. In order to understand if choices of analysis strategies (here: group comparison vs. correlative approach) may have caused this difference, we compared in a post-hoc analysis the 20 subjects with the highest ADHD Index with those 20 participants exhibiting the lowest score, in our sample. Also here, we found no significant differences in *global modularity Q* (*t* = 1.46; *p* = .15) or any of the other global measures reported above *(number of modules: t* = .71, *p* = .48; *average module size: t* = .60, *p* = .54; *variability in module size: t* = .67, *p* = .50).

To explore the possibility that alterations in *global modularity* may be restricted to clinically affected subjects (as observed, e.g., for autism: Rudie et al., 2013, or Alzheimer’s disease: De Haan et al., 2012), we conducted a further post-hoc analysis on the current data and compared *global modularity Q* values of the eight subjects with a clinical ADHD diagnosis with those of all other subjects. Also here, we did not observe a significant difference (Mann-Whitney U Test, one-tailed, *z* = .62, *p* = .27). However, this test relies on a comparison of 283 healthy subjects with only eight affected patients. Thus, we finally compared also the eight subjects with ADHD diagnosis with the 20 subjects of the first post-hoc analysis (20 persons with lowest ADHD Index). Also here, there was no significant difference (Mann-Whitney U Test, one-tailed, *z* = .08, *p* = .47). Similar results were obtained for all other global network measures *(number of modules: z* = 1.16., *p* = .12; *average module size: z* = .62, *p* = .27; *variability in module size: z* = .53, *p* = .58).

For comparability with studies that investigated the relationship between ADHD and functional connectivity strength within or between standard (i.e., group-average) brain networks (e.g., Sidlauskaite et al., 2016), we applied a canonical 400-node parcellation (Schaefer et al., 2018) to each individual’s functional MR scan and then assigned each node to one of seven well-established functional brain networks, i.e., visual, somato-motor, dorsal attention, ventral attention, limbic, frontoparietal, or default-mode network (Yeo et al., 2011). Examining the relationship of connectivity strength between/within these networks and ADHD-related behaviors, we observed no significant effects (Pearson correlation controlled for effects of age, sex, handedness, FSIQ, and mean FD; all *r* < .11, *p* > .056; Bonferroni corrected threshold: *p* = .0018; see Supplementary Table S3).

To assess the possibility that our estimates of functional connectivity (and thus also our graph-theoretical measures) may have been affected by distance-dependent motion artifacts which could remain in the data even after motion correction (Power et al., 2012; Ciric et al., 2017), we calculated for each edge the correlation between a) the association of its functional connectivity strength with mean framewise displacement, and b) the Euclidean distance between the two nodes of this respective edge (Ciric et al., 2018; note that for computational reasons this analysis is also based on the 400-node parcellation of Schaefer et al., 2018; see also above). As illustrated in Supplementary Figure S1, we observed no indications of distance-dependent artefacts. As further post-hoc control analysis addressing potential influences of head-motion, we tested whether there is a relationship between the ADHD index and the number of low-motion frames (FD<.2mm) in our sample. This was not the case (*r* =.01; p=.93). Nevertheless, we repeated all our analyses with the number of low-motion frames as covariate of no interest (rather than mean framewise displacement) and observed that the graph-theoretical results were almost unchanged, i.e., only slight changes were observed in respect to *p_i_* and *z_i_* (see Supplementary Table S5-7 and Supplementary Figure S2,3). Thus, these control analyses provide no evidence for influences of residual, distance-dependent motion artifacts on our results.

Lastly, to further characterize the functional role of the areas significantly associated with ADHD, i.e., we examined whether a) ADHD-related brain regions have generally higher *participation coefficient* than other regions of the brain (which would make them important as inter-module connectors in the sense of, e.g., the so-called diverse club; Bertolero et al., 2017) and b) whether ADHD-related brain regions have generally higher *within-module degree* than other regions of the brain, which would indicate a function as local hubs within their own modules. To this end, we extracted mean (group-average) values of *participation coefficient p_i_* and *within-module degree z_i_* from all significant clusters (Table 3,4) and determined their rank position within the whole-brain (group-average) distributions. In respect to *participation coefficient* all ADHD-associated regions scored around the center (rank positions between 20 and 80%), i.e., not in the extremes of the whole-brain *p_i_*-distribution (see Supplementary Table S4). Although most ADHD-related brain regions were also located around the center of the *z_i_*-distribution, we observed very high *z_i_* values (rank position > 80%) in middle frontal gyrus, anterior cingulate cortex, precuneus, and in both central clusters, and very low *z_i_* values (rank position < 20%) in posterior cingulate cortex, left posterior fusiform gyrus, and in both temporal clusters.

## Discussion

The current study investigated whether ADHD-related behaviors are associated with one of the key determinants of human brain function, i.e., brain network modularity. In contrast to previous studies that relied on ADHD vs. control group comparisons, we here applied a correlative approach investigating the appearance of ADHD symptoms across a broad range of variation. These ADHD-related behaviors correlated with region-specific but not global aspects of modularity, consistent with neurocognitive models of ADHD relating intrinsic connectivity between functional brain networks to ADHD. Our results extend these models to the non-clinical range of attention and executive functions.

### No association between ADHD symptoms and global modularity

The *default-mode interference hypothesis* (Sonuga-Barke & Castellanos, 2007) postulates stronger connectivity between the default-mode network and task-positive regions, i.e., a shift towards more integration and less segregation between these networks. This assumption was recently supported by two group-comparison studies that observed higher connectivity between default-mode network and task-positive regions in ADHD patients during cognitive tasks (Mills et al., 2018; Mowinckel et al., 2017). We found no direct support for this assumption in our non-clinical sample (post-hoc analysis; functional connectivity between standard networks). The graph-theoretical measure of global modularity considers simultaneously all connections within the entire network and indicates the general level of network segregation. Changes in global modularity can therefore occur in the presence of stronger or weaker connectivity between particular networks. Lin and colleagues (2014) reported higher global modularity of intrinsic functional brain networks in ADHD. This result, however, could not be replicated by Barttfeld and colleagues (2014), and further evidence from clinical ADHD samples is currently lacking. In our study focusing on behavioral variation across a broad non-clinical range, we also did not find support for an association with *global modularity*. However, despite the relatively large sample size, our data cannot be considered as strong evidence (in terms of Bayes Factors) against the presence of such associations.

While previous studies reported significant alterations in *global modularity* in patients with psychiatric conditions (Rudie et al., 2013; De Haan et al., 2012), it is still unclear whether global modularity relate to individual differences in cognitive abilities in the unimpaired brain (Stevens et al., 2012; Liang et al., 2016). We therefore speculated previously (Hilger et al., 2017b) that differences in modular network organization might become pronounced at a global level only in persons with severe neurological or psychiatric diseases. However, a post-hoc analysis on the current data (albeit underpowered) suggests that the eight ADHD-diagnosed subjects in the present sample did not differ in terms of *global modularity* from subjects without diagnosis.

### Region-specific connectivity profiles are associated with ADHD symptoms

In general, modular brain networks are organized in a way that balances functional integration and functional segregation (Gallos et al., 2012). Whereas the coordination and integration of different cognitive processes has been suggested to rely on exchange of information between different modules (high *participation coefficient*), the effectiveness of specific cognitive functions may be supported by less diverse, more focused processing of information within only one or between only few circumscribed modules (low *participation coefficient*; Bertolero et al., 2015, 2017; Gratton et al., 2012; Warren et al., 2014).

High within-module connectivity reflects that a node or brain region has strong influence on (or is highly influenced by) other nodes within the same functional module, and is therefore thought to facilitate more segregated specific cognitive functions (Warren et al., 2014; Gratton et al., 2012). In contrast, low values reflect less influence and more flexible (or independent) coupling to nodes within their modules. We observed region-specific patterns of positive and negative associations between non-clinical ADHD symptoms and within-module degree, suggesting that not only between-module interactions but also the quality of information flow within specific modules may be relevant for ADHD. Recent empirical evidence suggests further that both higher and lower levels of integration or segregation are important for cognitive performance (Cohen & D’Esposito, 2016; Hilger et al., 2017b). Our results support this and provide converging evidence from the domain of ADHD-associated behaviors. However, our outcome measure spans a wide range of behaviors and cognitive attitudes from impulsivity to self-esteem, and thus lacks the specificity needed to relate specific cognitive sub-functions to specific patterns of connectivity.

Nine mostly bilaterally located brain regions demonstrated significant effects in both *participation coefficient* and *within-module degree*. Interestingly, these associations were of opposite directions in all cases, i.e., high *p_i_*/low *z_i_*, or vice versa. This may indicate that in persons with higher ADHD Index the connectivity profile of these regions may be biased towards one type of information flow (distributed across modules or focused within modules). ADHD-associated regions, however, do not seem to have particular node-properties (post-hoc analysis). The mechanisms linking individual variations in modular brain network organization and subclinical variations in attention and executive functioning are thus not localized to particularly integrative (members of the diverse club; Bertolero et al., 2017) or particularly locally central regions.

### Partial support for network models of ADHD

ADHD is a complex phenomenon, involving atypical neural activation in distributed brain regions (Cortese et al., 2012; Dickstein et al., 2006), dysfunction of specific neural networks (Sidlauskaite et al., 2016; Konrad & Eickhoff, 2010), and fundamental alterations in intrinsic connectivity (Zhang et al., 2016; Di Martinos et al., 2013). To reiterate, both two- and three-network theories specifically suggest altered connectivity between the default-mode and task-positive brain networks (Sonuga-Barke & Castellanos, 2007; Cortese at al., 2012). At the most general level, our results support these network-based models by demonstrating significant and systematic relationships between functional brain network organization and variations in ADHD-related behaviors. General support for existing models also comes from our observation of localized rather than global effects, which is consistent with the focus on specific inter-module connection patterns in the current literature (Sripada et al., 2014; Choi et al., 2013). Although we did not observe associations of between-module connectivity strength with ADHD when using a standard network partition (post-hoc analysis), in our main analyses (based on individual partitions) we found lower within-module connectivity in circumscribed clusters adjacent to classical default-mode networks in persons with higher ADHD Index. This is in line with previous studies with ADHD samples (e.g., Kessler et al., 2014; Castellanos et al., 2008; Sripada et al., 2014) and suggests that altered DMN connectivity profiles might also exist in subjects with non-clinical difficulties in attention and executive function. It is plausible to assume that this association is less pronounced within the healthy range, and the use of individualized network partitions might have helped to detect such covariations within our non-clinical sample. Future research will be required to clarify how this alteration of within-module connectivity may relate to the diminished suppression of DMN activation that was suggested as cause of distracting intrusions and attentional lapses in ADHD (Sonuga-Barke & Castellanos, 2007).

Our results can also be related to recent three-network theories of ADHD, which propose stronger interactions between salience network and default-mode network, relative to weaker interactions between salience network and central executive network (Choi et al., 2013, for review see Castellanos & Aoki, 2016). As we observed significant associations in regions associated with the default-mode, salience, and central executive networks, in general our results support three-network theories. Nonetheless, both metrics studied here allow no conclusions about directionality of these connections, e.g., to which brain regions the salience network is connected more strongly.

Importantly, the associations reported in the current study were observed across a broad and continuous range of non-clinical behavioral variations. They may thus represent more general mechanisms linking intrinsic network organization to variations in behaviors that in ‘extreme’ expressions are associated with ADHD. This supports continuous conceptualizations of ADHD (Marcus et al., 2012) and suggests that ADHD is not only the extreme end in terms of behavioral variations (Levy et al., 1997) but also in terms of biological variations.

### Limitations

The CAARS ADHD Index has a high validity (Kooij et al., 2008; Erhardt et al., 1999). Nevertheless, there was no perfect match between those participants with highest ADHD Index and those reporting a clinical diagnosis. Further, it has been demonstrated that different ADHD measures can lead to slightly different results (Kooij et al., 2008), so that the dependency on the predictor measure may be addressed by future research. A second limitation is the rather short duration of the resting-state scan. While similar scan lengths are common in current functional connectivity research and while it has been shown that even less than 2min of fMRI can be used to build robust individual connectotypes (Miranda-Dominguez et al., 2014), it has recently been demonstrated that short scan durations can lead to systematic biases in graph-theoretical measures, e.g., reduced global modularity estimates (Gordon et al., 2017). Our rather large dataset may compensate for this problem to a certain degree, but future work will have to replicate the present results in datasets with longer scan durations. Finally, even though resting-state connectivity supposedly reflects fundamental organizational principles of the human brain (Biswal et al., 1995), and functional connectivity during cognitive demands may rely on these intrinsic properties (Cole et al., 2014, 2016), we consider it an important issue for future research to investigate whether the same associations persist in the presence of cognitive tasks.

## Conclusion

We demonstrate that non-clinical variations in ADHD symptoms relate significantly to the modular organization of human functional brain networks. Even though ADHD-related behaviors seem to vary independent of *global modularity* differences, region-specific profiles of between-module and within-module connectivity covary with the self-rated presence of (non-clinical) ADHD symptoms. Our results support a network perspective of ADHD and suggest that intrinsic functional connections between and within neuronal systems are relevant for a comprehensive understanding of individual variations in ADHD-related cognition and behavior.

## Acknowledgements

The authors thank Caterina Gawrilow for valuable comments on an earlier version of this manuscript. The research leading to the reported results has received funding from the German Research Foundation (DFG grant n° FI 848/6-1) awarded to CJF. Data were provided by the Nathan S. Kline Institute for Psychiatric Research (NKI), founded and operated by the New York State office of mental health. Principal support for acquisition of the data used in this project was provided by the NIMH BRAINS R01MH094639-01 grant for Michael Milham. Funding for the decompression and augmentation of administrative and phenotypic protocols was provided by a grant from the Child Mind Institute (1FDN2012-1). Additional personnel support was provided by the Center for the Developing Brain at the Child Mind Institute, as well as NIMH R01MH081218, R01MH083246, and R21MH084126.

## Author contributions statement

K.H. conducted the analyses. K.H. and C.F. conceived of the study, interpreted the results, and wrote the manuscript.

## Data availability statement

The data used in the present work were made available by the 1000 Functional Connectomes Project INDI and can be accessed under the following link: http://fcon_1000.projects.nitrc.org/indi/enhanced/. The code used in the current study has been deposited on github at https://github.com/KirstenHilger/ADHD-Modularity (https://doi.org/10.5281/zenodo.2574588).

