## Supplementary material for "ADHD Symptoms are Associated with the Modular Structure of Intrinsic Brain Networks in a Representative Sample of Healthy Adults": SI_Tables_Figures

* Corresponding author:

Dr. Kirsten Hilger
Goethe University
Department of Psychology
Theodor-W.-Adorno-Platz 6, PEG
D-60323 Frankfurt am Main

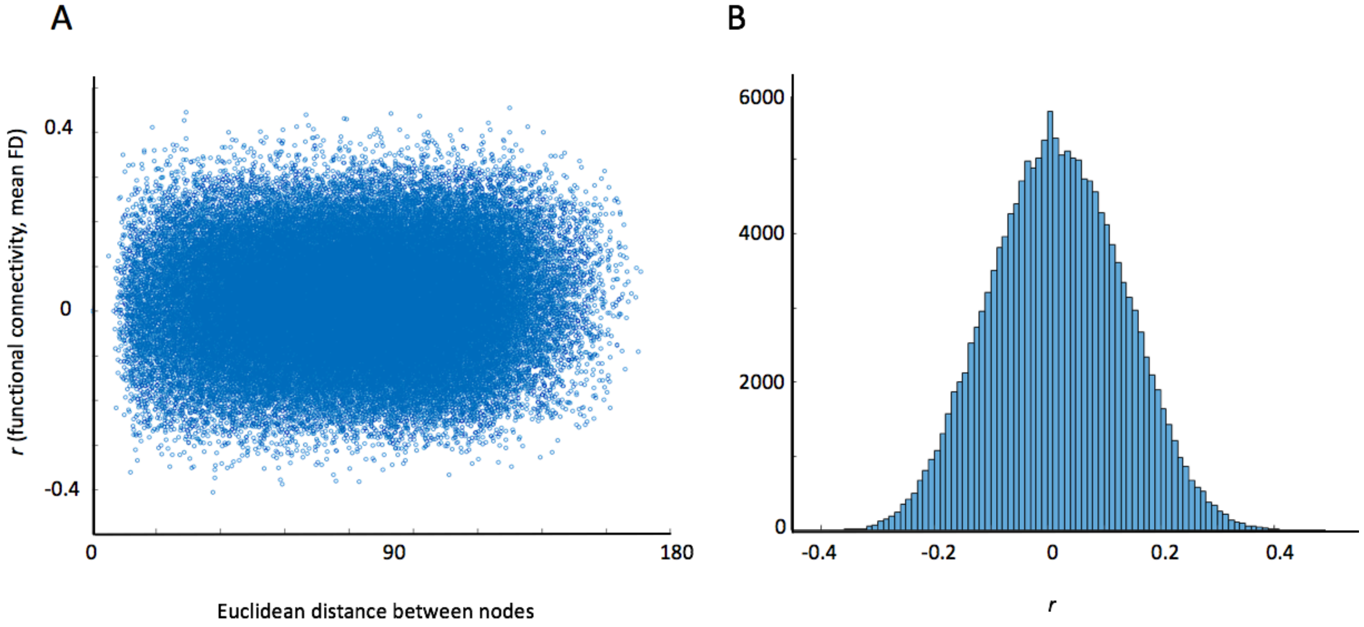


**Supplementary Figure S1.** Check for distance-dependent influences of in-scanner head motion on functional connectivity values. A, Scatterplot illustrating the correlation between each edge's functional connectivity strength and mean framewise displacement (*y*-axis) in dependency of Euclidean distance between the respective nodes of this edge (in mm, *x*-axis). B, Histogram of the correlation scores for the association between mean framewise displacement and functional connectivity values. FD, mean framewise displacement; *r*, Pearson correlation. Functional connectivity values are based on the 400-node parcellation of Schaefer et al. (2018).

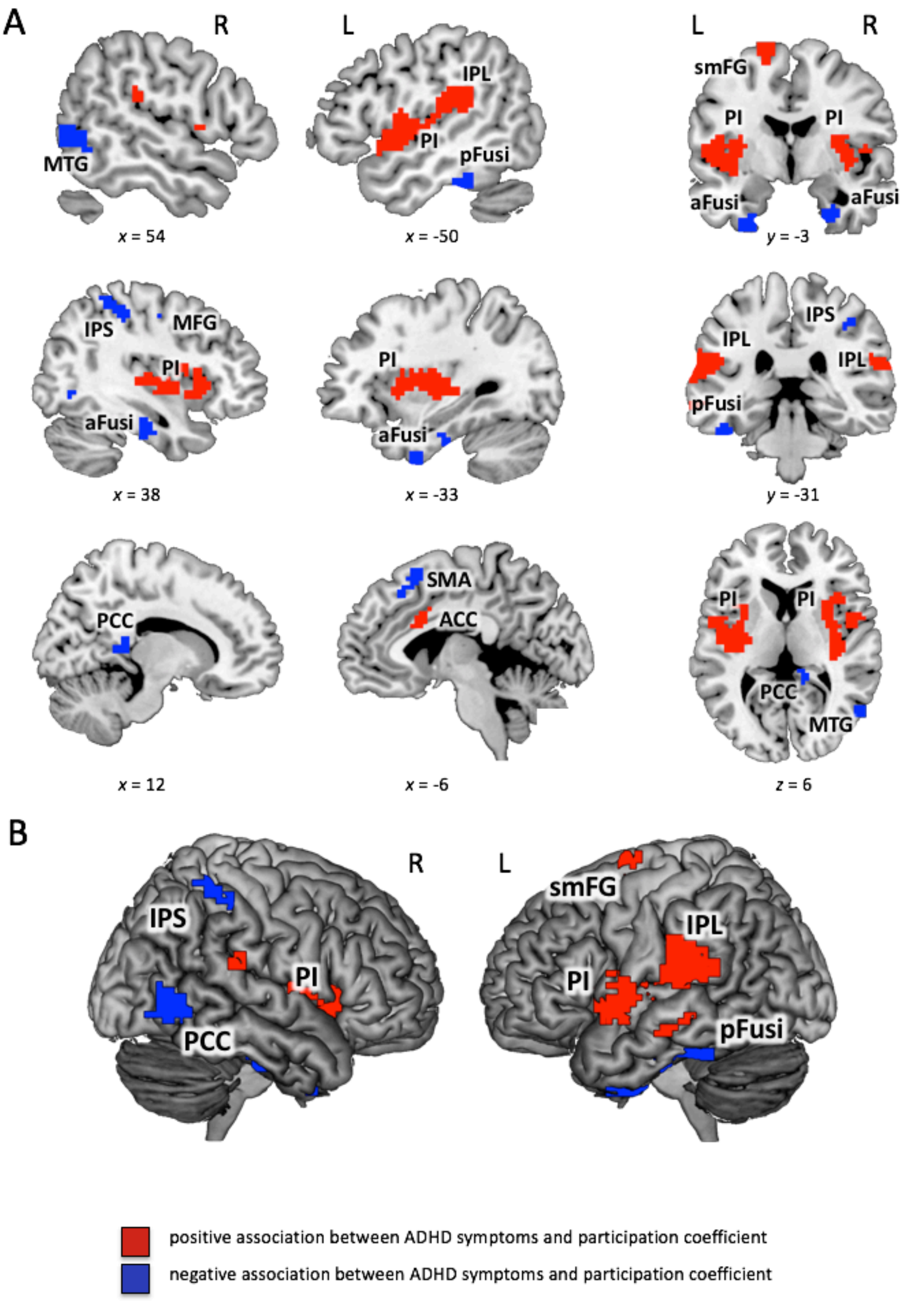


**Supplementary Figure S2.** Significant associations between Conners' ADHD Index and participation coefficient controlled for number of low-motion frames (rather than mean framewise displacement; see Post-Hoc Analyses in the Results Section of the Main Text, and Supplementary Table S6). *Participation coefficient* $p_i$ (see Methods for details) was calculated for binarized and proportionally thresholded graphs using five thresholds (graphs were defined by the top 10%, 15%, 20%, 25%, or 30% of strongest edges). Input for analyses were the individual mean maps for *participation coefficient* $p_i$, which were calculated by averaging across these five thresholds for each participant separately. Statistic parametric maps of *participation coefficient* $p_i$ are shown at a voxel-level threshold of $p < .005$ (uncorrected) combined with a cluster-level threshold of $k > 26$ voxels, corresponding to an overall family-wise error corrected threshold of $p < .05$ (see Methods). **(A)** Slice view; the *x*-, *y*-, and *z*-coordinates represent coordinates of the Montreal Neurological Institute template brain (MNI152).

**(B)** Render view; projection to the surface of the brain, search depth 12 voxels. PI, posterior insula; IPL, inferior parietal lobe; IPS, intraparietal sulcus; ACC, anterior cingulate cortex; MFG, middle frontal gyrus; SMA, supplementary motor area; aFusi, anterior fusiform gyrus; pFusi, posterior fusiform gyrus; PCC, posterior cingulate cortex; MTG, middle temporal gyrus; smFG, superior medial frontal gyrus.

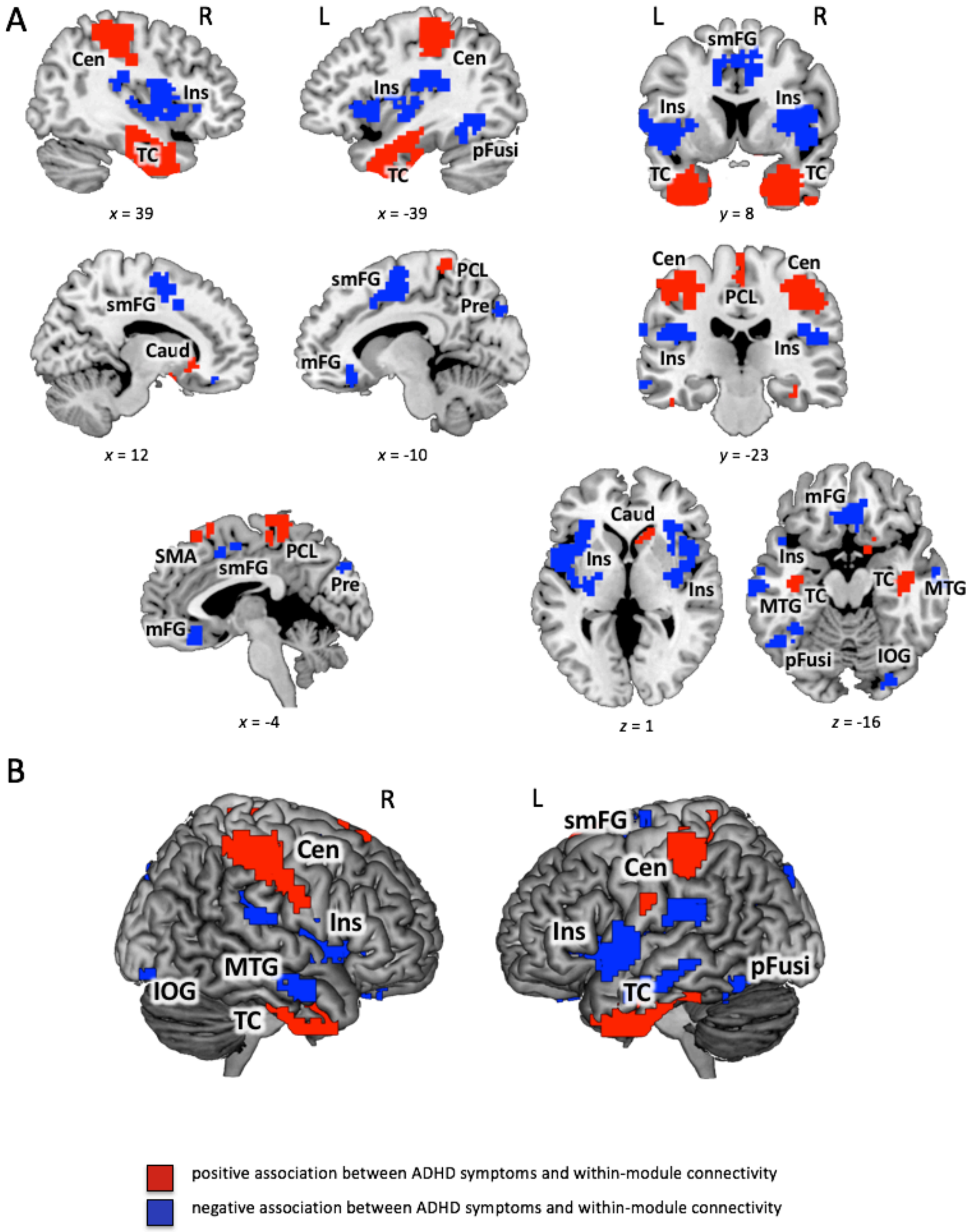


**Supplementary Figure S3.** Significant associations between Conners' ADHD Index and within-module degree controlled for number of low-motion frames (rather than mean framewise displacement; see Post-Hoc Analyses in the Results Section of the Main Text, and Supplementary Table S7). *Within-module degree* $z_i$ (see Methods for details) was calculated for binarized and proportionally thresholded graphs using five thresholds (graphs were defined by the top 10%, 15%, 20%, 25%, or 30% of strongest edges). Input for analyses was the individual mean maps for *within-module degree* $z_i$, which were calculated by averaging across these five thresholds for each participant separately. Statistic parametric maps of *within-module degree* $z_i$ are shown at a voxel-level threshold of $p < .005$ (uncorrected) combined with a cluster-level threshold of $k > 26$ voxels, corresponding to an overall family-wise error corrected threshold of $p < .05$ (see Methods). **(A)** Slice view; the *x*-, *y*-, and *z*-coordinates represent coordinates of the Montreal Neurological Institute template brain (MNI152). **(B)** Render view; projection to the surface of the brain, search depth 12 voxel. TC, temporal cluster comprising also amygdala, hippocampus, and parts of fusiform gyrus; Cen, central cluster spreading across central and postcentral sulci from precentral gyri and postcentral gyri to the inferior parietal lobes (comprising supramarginal gyri and anterior parts of intraparietal sulci); PCL, paracentral lobule; mFG, medial frontal gyrus; Ins, insular cluster comprising also parts of putamen, superior temporal gyrus, inferior frontal gyrus and inferior parietal lobe; MTG, middle temporal gyrus; pFusi, posterior fusiform gyrus; Pre, precuneus; IOG, inferior occipital gyrus; smFG, superior medial frontal gyrus; SMA, supplementary motor area; Caud, caudate nucleus.

**Supplementary Table S1.** Correlations between CAARS subscales and whole-brain characteristics of modular network organization

| | $r_{part.}$ | $p_{part.}$ |
|---|---|---|
| *Inattention/Memory Problems* | | |
| global modularity | .051 | .387 |
| number of modules | -.050 | .400 |
| average module size | .043 | .471 |
| variability in module size | -.033 | .579 |
| *Hyperactivity/Restlessness* | | |
| global modularity | .031 | .596 |
| number of modules | .007 | .912 |
| average module size | -.024 | .681 |
| variability in module size | -.010 | .863 |
| *Impulsivity/Emotional Lability* | | |
| global modularity | .082 | .166 |
| number of modules | .037 | .537 |
| average module size | -.055 | .356 |
| variability in module size | -.019 | .747 |
| *Self-Concept Problems* | | |
| global modularity | .147 | .013 |
| number of modules | -.062 | .300 |
| average module size | .061 | .307 |
| variability in module size | .118 | .046 |

$r_{part.}$, Pearson's correlation coefficient for the partial correlation controlling for effects of age, sex, handedness, mean framewise displacement, and Full Scale Intelligence Quotient; $p_{part.}$, *p*-value of significance for the partial-correlation.

**Supplementary Table S2.** Correlations between CAARS subscales and proportions of node type within the whole brain

| | $r_{part.}$ | $p_{part.}$ |
|---|---|---|
| *Inattention/Memory Problems* | | |
| ultra-peripheral nodes | .053 | .375 |
| peripheral nodes | .024 | .686 |
| non-hub connector nodes | -.019 | .751 |
| non-hub kinless nodes | -.07 | .239 |
| provincial hubs | .025 | .675 |
| connector hubs | .009 | .886 |
| kinless hubs | -.096 | .107 |
| *Hyperactivity/Restlessness* | | |
| ultra-peripheral nodes | .061 | .307 |
| peripheral nodes | -.026 | .658 |
| non-hub connector nodes | .010 | .871 |
| non-hub kinless nodes | -.006 | .916 |
| provincial hubs | -.042 | .480 |
| connector hubs | .048 | .418 |
| kinless hubs | -.074 | .213 |
| *Impulsivity/Emotional Lability* | | |
| ultra-peripheral nodes | .115 | .052 |
| peripheral nodes | -.038 | .518 |
| non-hub connector nodes | .025 | .675 |
| non-hub kinless nodes | -.038 | .52 |
| provincial hubs | -.015 | .800 |
| connector hubs | .003 | .956 |
| kinless hubs | -.063 | .285 |
| *Self-Concept Problems* | | |
| ultra-peripheral nodes | .042 | .480 |
| peripheral nodes | .068 | .251 |
| non-hub connector nodes | -.127 | .032 |
| non-hub kinless nodes | -.080 | .180 |
| provincial hubs | .015 | .802 |
| connector hubs | -.026 | .656 |
| kinless hubs | .034 | .570 |

$r_{part.}$, Pearson's correlation coefficient for the partial correlation controlling for effects of age, sex, handedness, mean framewise displacement, and Full Scale Intelligence Quotient; $p_{part.}$, *p*-value of significance for the partial-correlation.

**Supplementary Table S3.** Correlations between Conner's ADHD Index and functional connectivity strength within/between canonical brain networks

| | **VIS** | **SOM** | **DAN** | **VAN** | **LIM** | **FPN** | **DMN** |
|---|---|---|---|---|---|---|---|
| **VIS** | .011<br>(.847) | - | - | - | - | - | - |
| **SOM** | .012<br>(.833) | .007<br>(.901) | - | - | - | - | - |
| **DAN** | .016<br>(.782) | .016<br>(.784) | .021<br>(.717) | - | - | - | - |
| **VAN** | .035<br>(.552) | .024<br>(.684) | .107<br>(.069) | .031<br>(.591) | - | - | - |
| **LIM** | .024<br>(.675) | .004<br>(.939) | .043<br>(.463) | .084<br>(.635) | -.010<br>(.866) | - | - |
| **FPN** | -.015<br>(.794) | -.011<br>(.848) | .028<br>(.635) | .050<br>(.395) | -.022<br>(-.702) | .034<br>(.561) | - |
| **DMN** | -.086<br>(.142) | -.092<br>(.118) | -.102<br>(.084) | -.061<br>(.304) | -.044<br>(.449) | -.017<br>(.775) | .113<br>(.056) |

Pearson's correlation coefficient for the partial correlation between functional connectivity strength and ADHD symptoms controlling for effects of age, sex, handedness, mean framewise displacement, and Full Scale Intelligence Quotient; corresponding *p*-values are depicted in brackets; VIS, visual network; SOM, somato-motor network, DAN, dorsal attention network, VAN, ventral attention network, LIM, limbic network, FPN, frontoparietal network, DMN, default-mode network (Yeo et al., 2011).

**Supplementary Table S4.** Rank position of ADHD symptoms-related brain regions, relative to the whole-brain distribution of *participation coefficient* $p_i$ and *within-module degree* $z_i$

| Brain Region | BA | Hem | rank position $p_i$ | rank position $z_i$ |
|---|---|---|---|---|
| *positive association with* $p_i$ | | | | |
| posterior insula* | 13 | L | 68.66 | 79.43 |
| posterior insula, putamen* | 13 | R | 70.97 | 79.92 |
| anterior cingulate cortex | 24 | L | 61.91 | 50.86 |
| superior medial frontal gyrus* | 6 | L | 43.39 | 42.40 |
| inferior parietal lobe* | 40 | L | 50.75 | 74.80 |
| *negative association with* $p_i$ | | | | |
| anterior cingulate cortex | 32, 9 | L | 43.18 | **91.12** |
| middle frontal gyrus | 6 | R | 31.69 | **83.95** |
| supplementary motor area | 8, 6 | L | 66.32 | 24.55 |
| posterior fusiform gyrus | 20, 36 | L | 43.03 | **4.17** |
| intraparietal sulcus* | 40 | R | 47.94 | 78.98 |
| posterior cingulate cortex | | R | 64.93 | **1.31** |
| middle temporal gyrus | 37, 19 | R | 44.64 | 52.68 |
| inferior parietal lobe | 40 | R | 35.64 | 40.59 |
| *positive association with* $z_i$ | | | | |
| supplementary motor area | 8, 6 | R/L | 33.31 | **12.45** |
| temporal cortex, amygdala, hippocampus, fusiform gyrus | 38, 20, 28 | R | 52.29 | **12.64** |
| temporal cortex, amygdala, hippocampus, fusiform gyrus | 20, 38 | L | 49.09 | **12.38** |
| precentral gyrus, postcentral gyrus, inferior parietal lobe* | 3, 40 | R | 40.72 | **83.64** |
| precentral gyrus, postcentral gyrus, inferior parietal lobe | 3, 40 | L | 37.86 | **82.82** |
| paracentral lobule | 6, 4 | L | 38.69 | 23.32 |
| *negative association with* $z_i$ | | | | |
| medial frontal gyrus | 11, 32, 25 | R | 50.21 | 41.27 |

| | | | | |
|---|---|---|---|---|
| anterior cingulate cortex | 24 | R | 60.92 | 57.89 |
| insula, putamen, superior temporal gyrus, inferior frontal gyrus, inferior parietal lobule* | 13, 22, 40 | L | 57.35 | 71.16 |
| insula, putamen, superior temporal gyrus, inferior frontal gyrus, inferior parietal lobule* | 13, 47, 22 | R | 56.58 | 67.01 |
| superior medial frontal gyrus* | 6, 32, 24 | L | 52.13 | 77.52 |
| midde temporal gyrus | 21, 20 | R | 30.66 | 78.60 |
| thalamus | | R | 57.32 | 25.14 |
| thalamus | | L | 36.58 | 27.35 |
| posterior fusiform gyrus | 37, 19 | L | 59.31 | **17.80** |
| posterior fusiform gyrus | | R | 35.77 | 26.18 |
| posterior cingulate cortex | 30 | L | 57.75 | 66.01 |
| precuneus | 19, 7, 31 | R/L | 53.49 | **81.13** |
| inferior occipital gyrus | 18 | R | 39.84 | 36.88 |

BA, approximate Brodmann's area; Hem, hemisphere; L, left; R, right; regions with significant effects in both measures (participation coefficient and within-module degree) are marked with an asterisk and separately listed in Table 5; rank positions < 20% or > 80% are depicted in bold letters.

**Supplementary Table S5.** ADHD symptoms and global modularity measures controlled for number of low-motion frames (rather than mean framewise displacement; see Post-Hoc Analyses in the Results Section of the Main Text)

| | $r_{part.}$ | $p_{part.}$ | **$BF_{01}$-Reg.** |
|---|---|---|---|
| *Whole-brain modularity measures* | | | |
| global modularity | .11 | .059 | 0.63 |
| number of modules | -.09 | .148 | 1.35 |
| average module size | .08 | .204 | 1.89 |
| variability in module size | -.03 | .626 | 3.13 |
| *Whole-brain proportions of node types* | | | |
| ultra-peripheral nodes | .01 | .824 | 3.80 |
| peripheral nodes | .06 | .359 | 2.27 |
| non-hub connector nodes | -.07 | .220 | 2.28 |
| non-hub kinless nodes | -.10 | .107 | 1.02 |
| provincial hubs | .09 | .149 | 1.33 |
| connector hubs | -.07 | .253 | 2.57 |
| kinless hubs | -.07 | .263 | 2.61 |

$r_{part.}$, Pearson's correlation coefficient for the partial correlation controlling for effects of age, sex, handedness, number of low-motion frames (FD < 0.2mm), and FSIQ; $p_{part.}$, *p*-value of significance for the partial-correlation; $BF_{01}$-Reg., Bayes Factor in favor of the null hypothesis (i.e., absence of correlation). Bayes Factors were calculated for linear regression models predicting ADHD Index values by the respective whole-brain measure of modular network organization or whole-brain proportions of node types, respectively, while effects of age, sex, handedness, number of low-motion frames (FD < 0.2mm), and FSIQ were controlled.

**Supplementary Table S6.** ADHD symptoms and participation coefficient controlled for age, sex, handedness, FSIQ, and number of low-motion frames (rather than mean framewise displacement, see Post-Hoc Analyses in the Results Section of the Main Text)

| Brain Region | BA | Hem | x | y | z | $t_{max}$ | *k* |
|---|---|---|---|---|---|---|---|
| *positive association* | | | | | | | |
| posterior insula, inferior parietal lobe | 13, 40 | L | -57 | -36 | 21 | 5.45 | 840 |
| posterior inula, putamen | 13 | R | 36 | -9 | 3 | 3.64 | 384 |
| anterior cingulate cortex | 24 | L | -6 | 18 | 24 | 3.60 | 110 |
| superior medial frontal gyrus | 6 | L | -18 | -3 | 69 | 3.32 | 47 |
| inferior parietal lobe | 40 | R | 63 | -30 | 21 | 3.18 | 33 |
| *negative association* | | | | | | | |
| middle frontal gyrus | 6 | R | 33 | -12 | 45 | 2.97 | 31 |
| supplementary motor area | 8, 6 | L | -6 | 18 | 57 | 3.87 | 40 |
| anterior fusiform gyrus | 28, 38 | R | 27 | 6 | -39 | 3.34 | 79 |
| anterior fusiform gyrus | 20, 36 | L | -30 | 3 | -39 | 3.25 | 63 |
| posterior fusiform gyrus | 20, 37 | L | -48 | -33 | -27 | 4.09 | 77 |
| posterior fusiform gyrus | 20 | R | 42 | -18 | -27 | 3.70 | 46 |
| intraparietal sulcus | 40 | R | 33 | -36 | 48 | 3.38 | 91 |
| posterior cingulate cortex | | R | 12 | -39 | 9 | 3.84 | 27 |
| middle temporal gyrus | 37, 19 | R | 57 | -63 | 0 | 3.71 | 105 |

BA, approximate Brodmann's area; Hem, hemisphere; L, left; R, right; coordinates refer to the Montreal Neurological Institute template brain (MNI); $t_{max}$, maximum *t* statistic in the cluster; *k*, cluster size in voxels of size 3 x 3 x 3 mm.

**Supplementary Table S7.** ADHD symptoms and within-module degree controlled for age, sex, handedness, FSIQ, and number of low-motion frames (rather than mean framewise displacement, see Post-Hoc Analyses in the Results Section of the Main Text)

| **Brain Region** | **BA** | **Hem** | **x** | **y** | **z** | $t_{max}$ | ***k*** |
|---|---|---|---|---|---|---|---|
| *positive association* | | | | | | | |
| supplementary motor area | 8, 6 | R/L | 0 | 30 | 60 | 3.20 | 42 |
| caudate | | R | 15 | 24 | -6 | 3.90 | 39 |
| temporal cortex, amygdala, hippocampus, fusiform gyrus | 38, 20, 28 | R | 33 | 6 | -30 | 5.76 | 794 |
| temporal cortex, amygdala, hippocampus, fusiform gyrus | 20, 38 | L | -27 | -15 | -33 | 5.15 | 872 |
| precentral gyrus, postcentral gyrus, inferior parietal lobe | 3, 40 | R | 39 | -33 | 51 | 6.72 | 588 |
| precentral gyrus, postcentral gyrus, inferior parietal lobe | 3, 40 | L | -45 | -33 | 51 | 5.37 | 373 |
| paracentral lobule | 6, 4 | L/R | 0 | -33 | 72 | 3.88 | 180 |
| *negative association* | | | | | | | |
| medial frontal gyrus | 11, 32, 25 | R | 3 | 30 | -15 | 4.42 | 164 |
| insula, putamen, superior temporal gyrus, inferior frontal gyrus, inferior parietal lobule | 13, 22, 40 | L | -48 | 9 | -3 | 5.64 | 883 |
| insula, putamen, superior temporal gyrus, inferior frontal gyrus, inferior parietal lobule | 13, 47, 22 | R | 39 | 3 | 15 | 5.46 | 620 |
| superior medial frontal gyrus | 6, 32, 24 | L | -12 | -3 | 63 | 5.04 | 507 |
| midde temporal gyrus | 21, 20 | R | 63 | -6 | -21 | 4.48 | 63 |
| midde temporal gyrus | 21, 20 | L | -66 | -30 | -6 | 3.78 | 104 |
| cuneus/precuneus | 19 | R | 42 | -24 | 24 | 3.86 | 168 |
| posterior fusiform gyrus | 37, 19 | L | -36 | -48 | -12 | 4.26 | 139 |
| posterior cingulate cortex | 30 | L | -24 | -66 | 21 | 3.60 | 27 |
| precuneus | 19, 7, 31 | R/L | -12 | -87 | 36 | 3.84 | 85 |
| inferior occipital gyrus | 18 | R | 27 | -84 | -12 | 3.31 | 30 |

BA, approximate Brodmann's area; Hem, hemisphere; L, left; R, right. Coordinates refer to the Montreal Neurological Institute template brain (MNI); $t_{max}$, maximum $t$ statistic in the cluster; $k$, cluster size in voxels of size 3 x 3 x 3 mm.
